# dSTORMQuant: A Python Package for Post-Processing and Quantitative Analysis of SMLM datasets

**DOI:** 10.64898/2026.06.30.735216

**Authors:** Suraj Karki, Britta Nemeita, Anne Sophie Hammann, Sven Thoms

## Abstract

**Summary:** Single-molecule localization microscopy techniques, such as (direct) stochastic optical reconstruction microscopy ((d)STORM) and photo-activated localization microscopy (PALM) enable the visualization of subcellular molecular organization beyond the diffraction limit of conventional light microscopy. Not only is data acquisition rather slow, but the downstream analysis of localization datasets often remains computationally challenging and time-consuming. Consequently, the complexity and duration of data processing often limit experiments to the acquisition and analysis of only small numbers of cells or regions of interest, thereby restricting the statistical power and biological reliability of SMLM studies. To address this limitation, we developed an open-source Python-based package for automated, high-throughput post-processing and quantitative analysis of SMLM localization data, enabling efficient and straightforward handling of extensive datasets with minimal manual intervention.

**Availability and implementation:** dSTORMQuant (source code and documentation) are freely available on GitHub at https://github.com/BCMM-Bielefeld-University/dSTORMQuant under GPL v3 license.

## Introduction

With SMLM techniques, such as dSTORM (Heilemann et al. 2008), biological structures and molecular organization can be visualized with ∼20 nm resolution, thereby overcoming the resolution limit of conventional microscopy (Lelek et al. 2021). These methods rely on the stochastic collection of fluorescent molecules over a period of time, whose localizations are reconstructed into super-resolution images. As SMLM data fundamentally differ from conventional pixel-based imaging, specialized software is required for efficient filtering, processing, and analysis of localization datasets.

The raw lists of localizations generated during dSTORM imaging often contain millions of individual molecular coordinates, necessitating efficient and scalable processing workflows for extraction of biologically relevant information. Several open-source solutions for SMLM-analysis have been developed, including the FIJI-Plugin ThunderSTORM (Ovesný et al. 2014), MATLAB-based Grafeo (Haas and Peaucelle 2021), LOCAN (Doose 2022), and CODI, a cloud-based software which is associated with the ONI Nanoimager Hardware.

While these tools provide powerful functionalities for localization analysis, they typically require manual, file-by-file execution and do not integrate post-processing and downstream analysis steps within a single framework. For instance, tools such as ThunderSTORM require the user to process each dataset individually, with subsequent analysis steps, such as clustering, needing to be performed separately in additional software. As a result, these tools are not optimized for automated, high-throughput processing of large numbers of multi-color dSTORM localization datasets.

Here, we present a fully automated and scalable analysis workflow that combines multiple state-of-the-art algorithms and open-source approaches into a unified processing workflow for 2D-SMLM data. Our workflow enables automated batch processing of localization lists without the need for manual intervention or user-dependent stepwise execution. By integrating post-processing, quantitative analysis, and image reconstruction within a single framework, dSTORMQuant substantially improves throughput, reproducibility, and usability for large-scale SMLM studies. The software is primarily implemented in Python to ensure accessibility, interoperability, and straightforward integration across different computational environments.

## Methods

To enable standardized and scalable processing of dSTORM data acquired across different microscopy platforms, including the ONI Nanoimager, we developed a computational workflow supporting the simultaneous analysis of multiple localization tables.

The workflow accepts lists of single-molecule localizations as input, generated by NimOS (ONI Nanoimager) or other widely used localization software, such as rapidSTORM (Wolter et al. 2012).

Associated metadata, comprising file name, frame index, and channel assignment, are provided via a dedicated metadata table, while all processing and analysis parameters are centrally defined in a configuration file, allowing flexible adaptation to varying experimental and analytical requirements. For dSTORM data, an initial set of frames can be excluded from downstream analysis to discard localizations detected in a phase in which simultaneous excitation of large numbers of fluorophores occurs prior to the establishment of stable stochastic blinking conditions.

Our workflow includes drift correction, filtering, temporal grouping, and cell detection, with subsequent clustering, k-nearest neighbor (kNN), and colocalization analysis (Fig. 1A).

**Fig. 1:**
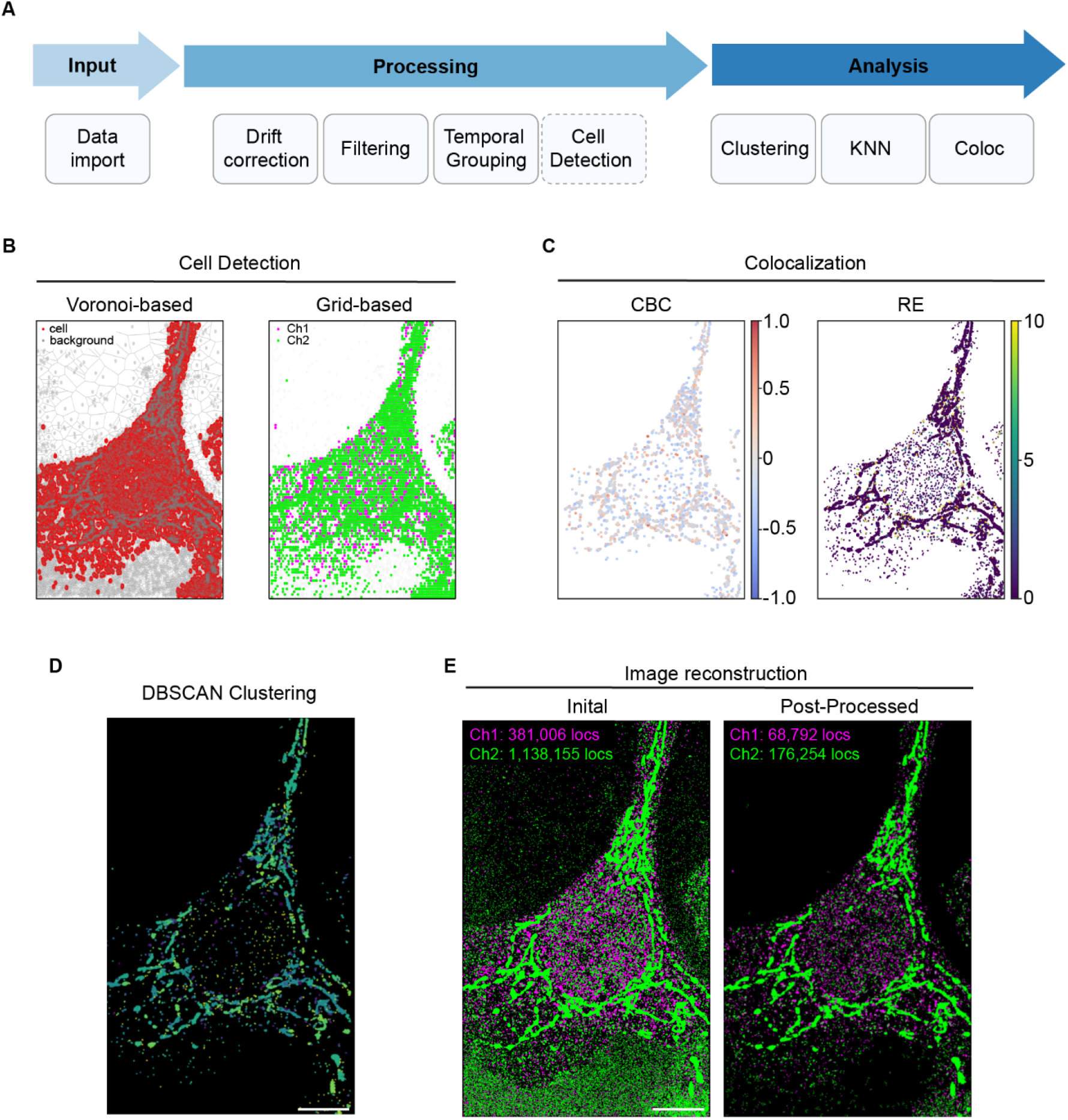
Overview of the dSTORMQuant analysis workflow. **A** Schematic illustration of the dSTORMQuant processing and analysis pipeline. Arrows indicate the sequential progression from input through processing to analysis. Individual modules are listed beneath each stage. The dashed border of the cell detection module denotes its optional use, depending on experimental requirements. **B-E** Example results of processing and analysis of a U2OS WT cell stained for mitochondria and peroxisomes as two different organelle markers: **B** Localization map of Voronoi- and grid-based cell detection. **C** Localization map by Coordinate-Based Co-localization (CBC) and Relative Enrichment (RE) methods. **D** DBSCAN (Density-based spatial clustering of applications with noise) result. Scale bar: 10 µm. **E** Reconstructed dSTORM image using the initial localization list in comparison with the post-processed localization list. The number of localizations per channel is noted on the upper left. Scale bar: 10 µm.

### Post-Processing

#### Drift correction

Mechanical instabilities of the microscope stage, thermal fluctuations, and external vibrations during data acquisition can cause systematic spatial displacements of localizations over time, a phenomenon referred to as drift. If left uncorrected, drift leads to blurring of the reconstructed super-resolution image and compromises the reliability of subsequent spatial analyses. We employed adaptive intersection maximization (AIM) (Ma et al. 2024) algorithm, a state-of-the-art approach for drift correction in single-molecule localization microscopy. Our implementation was adapted from the Picasso (Schnitzbauer et al. 2017) software package and extended to support CSV-based input/output handling, automated sanity checks, and HDF5 conversion for internal data processing.

To accommodate the sequential nature of two-channel acquisitions, as employed in dSTORM imaging, drift correction was applied independently to each channel, as each channel accumulates drift exclusively during its respective acquisition period. Following per-channel correction, the two channels were registered to a common coordinate system by subtracting the cumulative drift of channel 1 from all localizations of channel 2, thereby enabling accurate colocalization and inter-channel distance analyses.

To further ensure the integrity of the drift correction, a validation step was incorporated to prevent artifacts arising from sparse fluorophore blinking events toward the end of the acquisition. Under such conditions, AIM may operate on segments containing an insufficient number of localizations, potentially yielding spurious drift estimates. Segments for which the estimated drift exceeded a user-defined threshold were therefore identified as outliers, and their drift values were replaced by the mean drift computed over the preceding N valid segments.

### Filtering

Filters are applied step-by-step in a chosen order to remove low-quality localizations before performing analysis. Filtering criteria include the sigma (σ) of the point-spread function (PSF), photon count, localization precision, and p-value, enabling the exclusion of localizations with broad PSFs, low emission intensity, or poor spatial precision. The p-value filter provides an additional quality criterion derived from the statistical test performed during the fitting process. The stringency of filtering is fully user-controlled: by adjusting the respective threshold parameters, the user determines how strictly each criterion is applied and consequently how many localizations are retained or discarded. This allows the filtering workflow to be flexibly adapted to the specific quality requirements of a given dataset. Together, these filters substantially improve the quality of the retained localization (e.g., for representative two-color dataset shown in Fig. 1 the median localization precision is reduced from 13.5 nm to 9.6 nm for Channel 1 and 16.3 nm to 11.2 nm for Channel 2).

### Temporal Grouping

A fundamental challenge in localization-based super-resolution microscopy is overcounting, arising from fluorophores undergoing multiple blinking events that are localized across consecutive frames (Van De Linde and Sauer 2014). Such duplicate localizations can compromise both image quality and downstream quantitative analysis. To address this, dSTORMQuant applies temporal grouping, a spatiotemporal clustering procedure that merges localizations originating from the same blinking event based on fixed thresholds for spatial distance and frame separation (Ovesný et al. 2014). A cKDTree is constructed over all localization coordinates, and a breadth-first search identifies connected detections satisfying both spatial and temporal proximity criteria. All detections within a group are merged into a single representative localization using their mean spatial coordinates. Groups can additionally be filtered by temporal duration to exclude noise and artifacts, including anomalously short-or long-lived events.

### Cell detection

To confine analysis to cellular regions and minimize contributions from extracellular background, dSTORMQuant implements two localization density-based cell detection methods: grid-based (Mazouchi and Milstein 2016) and Voronoi-based cell detection (Voronoi 1908b, 1908a; Levet et al. 2015) (Fig. 1B). The grid-based approach subdivides the field of view into a user-defined 2D grid, counts localizations per bin, and classifies bins exceeding a defined density threshold as cellular, with remaining bins treated as background. For Voronoi-based detection, a Voronoi diagram is constructed from all localization coordinates, and a polygon area is derived for each region. Infinite and empty regions are discarded, and areas are clipped at a defined percentile to limit the influence of boundary effects. Otsu thresholding (Otsu 1979) is subsequently applied to partition localizations into cellular and background classes, with localizations whose Voronoi area falls at or below the threshold assigned to the cellular region.

### Quantitative Analysis

#### Nearest Neighbor Analysis

kNN is a well-established approach for quantifying spatial relationships between localizations for SMLM data. dSTORMQuant implements kNN-based spatial analysis at both the localization and cluster level, supporting intra-channel as well as inter-channel characterization. To accommodate diverse datasets and analytical requirements, the framework offers a selection of search algorithms and distance metrics for neighbor computation. For intra-channel analysis, the k-nearest neighbors of each localization within the same channel are identified, enabling the characterization of local molecular density and spatial organization. For inter-channel analysis, nearest-neighbor distances from each localization in a primary channel to its closest counterpart in a reference channel are computed for all pairs of distinct channels, providing a quantitative measure of molecular co-occurrence and spatial proximity. In addition, for both intra- and inter-channel analyses, the mean distance from each localization or cluster centroid to all neighbors within a fixed search radius r is calculated.

### Colocalization

To assess spatial association between distinct molecular species, dSTROMQuant incorporates two complementary colocalization methods: Coordinate-Based Co-localization (CBC) (Malkusch et al. 2012) and Relative Enrichment (RE) (Ejdrup et al. 2022) (Fig. 1C). CBC quantifies spatial overlap between two channels by calculating the fraction of neighboring localizations from the reference channel within a defined radius for each localization. This is evaluated bidirectionally and integrated following the implementation in LOCAN (Doose 2022). RE determines whether localizations of the primary channel are locally enriched or depleted relative to the reference channel, using Voronoi tessellation of the reference channel to define neighborhoods and comparing observed distributions against a random expectation.

### Clustering

Spatial clustering constitutes a fundamental component of SMLM analysis, enabling the identification of functionally relevant groups of localizations such as protein complexes or cellular organelles (Fig. 1D). Our package implements three clustering approaches, DBSCAN (Ester, Kriegel, and Xu 1996), HDBSCAN (Campello et al. 2015), and FINDER (Verzelli et al. 2022), which can be applied independently to the channels.

DBSCAN and HDBSCAN identify clusters of arbitrary shape based on local point density, with HDBSCAN extending this by extracting the most stable clusters across a density hierarchy while flagging outliers. FINDER offers a parameter-free alternative, automatically determining global clustering parameters to robustly detect molecular clusters while minimizing false positives and negatives in noisy and dense data; its use requires a C++ extension for Python. Localizations forming groups of fewer than three points are classified as noise. For each identified cluster, key properties, including localization count and density, as well as cluster centroid position and area, are recorded. Cluster centroids can subsequently serve as input for kNN analysis at the cluster level.

### Visualization and Output

After each processing step, the remaining localizations are saved to a new list, and histograms of all available localization parameters (background, intensity, localization precision, p-value, and sigma x and y) are exported as PNG files. Additionally, the applied drift correction values for x and y coordinates are saved. This output format facilitates direct data evaluation.

Super-resolution image reconstruction is performed using Napari (Sofroniew et al. 2026), providing an immediate visual assessment of the effect of each post-processing step on localization number and image quality (Fig. 1E). Display parameters can be further adjusted interactively after processing. For headless environments such as scientific cloud platforms, the Napari function can be disabled; image reconstruction of all datasets can then be performed locally at a later stage.

All quantitative analysis results of kNN, clustering, and colocalization are exported as CSV tables accompanied by corresponding PNG outputs, ensuring compatibility with downstream data handling workflows.

## Summary

With dSTORMQuant as a Python package, we have introduced a simple and straightforward way to process and analyze SMLM data. By combining multiple state-of-the-art metrics and algorithms with additional steps and modifications needed for 2D-dSTORM data, we have generated a workflow that is particularly easy to use and well-suited for data acquired with the ONI Nanoimager. However, the common input format allows for the processing and analysis of SMLM data, acquired with various microscopy hardware. The simultaneous analysis of multiple datasets enables quantitative evaluation of SMLM data not only within individual channels but also across channels, allowing researchers to investigate spatial relationships and interactions between different molecular species. Large datasets can be processed and analyzed more easily, eliminating the need to collect a small number of cells or images. This enables researchers to draw more reliable conclusions from their collected data.

## Supporting information

User guide

## Author contributions

Sven Thoms (Funding acquisition [lead], Supervision [lead], Writing – review & editing [equal]), Suraj Karki (Investigation [equal], Methodology [equal], Software [lead], Validation [equal], Writing – original draft [supporting], Writing – review & editing [equal]), Britta Nemeita (Investigation [equal], Methodology [equal], Validation [equal], Visualization [lead], Writing – original draft [lead], Writing – review & editing [equal]), Anne Sophie Hammann (Investigation [equal], Methodology [equal], Validation [equal], Writing – review & editing [equal])

## Supplementary material

The described Python package, including a detailed user guide, instructions for installation, and sample files, is freely available on GitHub at https://github.com/BCMM-Bielefeld-University/dSTORMQuant.

A User Guide for dSTORMQuant is available at *Bioinformatics* online.

## Acknowledgements

This work was supported by the Deutsche Forschungsgemeinschaft (TH1538/3-1 to ST), the Anschubfonds Medizinische Forschung (AMF) of Bielefeld University (projects AQP4x, RVtherapy, ERDKAR, and HC3P), de.NBI Cloud within the German Network for Bioinformatics Infrastructure (de.NBI) and ELIXIR-DE (Forschungszentrum Jülich and W-de.NBI-001, W-de.NBI-004, W-de.NBI-008, W-de.NBI-010, W-de.NBI-013, W-de.NBI-014, W-de.NBI-016, W-de.NBI-022), which provided computing resources to conduct the analysis. We gratefully acknowledge Marcel Müller for helpful discussions on this post-processing and analysis workflow. We thank Julia Hofhuis for testing dSTORMQuant. We thank Mark Schüttpelz for comments on this manuscript.

## Data availability

The data underlying this article are freely available at GitHub at https://github.com/BCMM-Bielefeld-University/dSTORMQuant.

