## Supplementary material for "dSTORMQuant: A Python Package for Post-Processing and Quantitative Analysis of SMLM datasets": User guide

Document version June 30th, 2026

**Abstract**

**Program Summary**

Program title: dSTORMQuant
Version: 0.0.1
Language: Python, C++;
Operating system: Any distribution (e.g. Linux, MacOSX, Windows);
License: GNU General Public License v3.0
Nature of problem: It consists of algorithms for post-processing and data analysis for SMLM datasets.

**Scientific Field**

Computational analysis of super-resolution microscopy data, focused on Single Molecule Localization Microscopy, including analysis of spatial molecule patterns, clustering of molecules, colocalization of different molecules, and nearest-neighbor distance statistics.

**Availability and Implementation**

dSTORMQuant is implemented primarily in Python with performance-critical clustering components in C++. It is distributed as open source and supports command-line execution with configurable YAML-based workflows. The source code is freely available from the GitHub Page: https://github.com/BCMM-Bielefeld-University/dSTORMQuant

**Keywords**

dSTORMQuant— SMLM— super-resolution microscopy— clustering— colocalization— drift correction— spatial

### 1. Introduction

dSTORMQuant is a modular toolbox of post-processing and analysis algorithms for super-resolution microscopy data such PALM, STORM, and dSTORM. It is primarily implemented in Python, with performance-critical components in C++ and optional Cython bindings. The project is published under the MIT License and is intended to make reproducible, configurable dSTORM data analysis accessible to researchers and developers. dSTORMQuant assumes localization coordinates with metadata as input and focuses on coordinate-based quantitative analysis rather than raw image fitting.

#### 1.1 What is dSTORMQuant?

Single-molecule localization experiments produce lists of detected emitters (x, y, frame, intensity, precision, etc.). These coordinate lists serve as the starting point for quantitative analysis that extends beyond image rendering. dSTORMQuant provides a pipeline and a collection of modular algorithms that operate on such lists to extract biologically and physically meaningful measures: localization precision estimates, spatial point-pattern statistics, cluster identification and characterization, temporal grouping of repeated detections, colocalization metrics, nearest-neighbor analyses, and cell-level segmentation/phenotyping. The codebase combines high-level Python orchestration with a C++ accelerated clustering implementation for larger datasets.

#### 1.2 What dSTORMQuant can do?

- Filter localization co-ordinates based on sigma, intensity, localization precision, and p-value.
- Correct experimental drift using the Adaptive Intersection Maximization (AIM) algorithm and segment-based validation.
- Merge temporally closed localization co-ordinates using temporal grouping.
- Detect cell boundary via grid or Voronoi approaches and extract per-cell metrics.
- Identify and characterize clusters using DBSCAN, HDBSCAN, and FINDER.
- Co-localization analysis using coordinate-based colocalization (CBC) and relative enrichment measures.
- Run nearest-neighbor analysis to characterize spatial organization, assess inter-channel relationships, cluster-level summaries, etc.
- Generate standardized quantitative outputs and visualization plots (via Napari-based integration and plotting utilities) suitable for downstream statistical analysis.
- Pipeline configuration and execution are controlled through YAML-based configuration (config.yaml) and command-line entry points.

##### 1.3 What dSTORMQuant cannot do?

dSTORMQuant is not a localization engine. It is designed for post-processing Single Molecule Localization Microscopy (SMLM) data and extracting quantitative information. It is also not a Graphical User Interface (GUI) application. Instead, it is a Python package that supports localization coordinates from STORM, dSTORM, and PALM datasets. dSTORMQuant emphasizes quantitative analysis of super-resolution microscopy data, including spatial point-pattern analysis, clustering, colocalization, and nearest-neighbor statistics.

### 2. Installation

#### 2.1 Requirements

- Python 3.12 or higher version
- Windows, Mac or Linux

#### 2.2 Installation

2.2.1 Clone git repository

- Install git (https://git-scm.com/) if needed. Use Terminal (Linux, MacOS) or Cmd (Win). Use **cd** to navigate to the target directory. (e.g. cd git)
- Type: git clone https://github.com/BCMM-Bielefeld-University/dSTORMQuant.git

2.2.2 Built the package

- Change directory: cd dSTORMQuant
- Build by entering command: pip install -e .

2.2.3 Build and install FINDER (if you want FINDER clustering algorithm)

- Change directory: cd finder_cpp
- Install requirements: pip install -r requirements.txt
- Build: pip install -e . --no-build-isolation
- Go back: cd ..

2.2.4 After successful built, you can check by entering: dSTORMQuant

#### 2.3 Available Scripts

The following CLI commands are supported when executed from the project directory.

| Command | Purpose |
|---|---|
| dSTORMQuant | Main entry command for the SMLM analysis pipeline. Runs the full processing workflow based on config/config.yaml. |
| merge-csv | Merge CSV outputs from multiple experiments into data/output/merged_channel_data/ grouped by cell line and channel. |
| save-napari-visualizations | Batch-save Napari views as images into each experiment's test_images/ folder. |

### 3. The Philosophy

All data processing steps in dSTORMQuant follows a staged workflow. First, the input file is validated, and the relevant metadata are loaded from the experiment. The localization coordinates are then partitioned into the appropriate channel-specific subsets according to the metadata and user-defined configuration. These subsets are processed independently through the selected analysis modules, including filtering, drift correction, temporal grouping, cell boundary detection, clustering, colocalization, and nearest-neighbor analysis. The complete cell can be analyzed and biologically quantified in one go.

Algorithm:

1. load configuration file C
2. load metadata table M and select the row for the current input file
3. load single-molecule localization list L
4. validate required columns and metadata fields
5. partition the localization list into channel-specific streams using the metadata-defined frame and channel ranges
6. Apply drift correction to the channel data
7. Apply filtering according to the configured threshold parameters
8. apply temporal grouping, if enabled
9. apply cell boundary detection, if enabled
10. compute nearest-neighbor statistics, if enabled
11. Run clustering, if enabled
12. compute colocalization metrics, if enabled
13. generate visual plots and summary plots
14. save the processed localization data and result files
15. archive the experiment outputs into standardized final results

#### 3.1 The Input File Format

For compatibility and consistency across experimental setups, dSTORMQuant adopts a structured CSV-based localization format as its standard input. The format is designed to accommodate the typical output of modern SMLM acquisition and localization software, including CODI, and covers the essential observables required for spatial, temporal, and photometric analysis.

Each input file is a comma-separated table in which every row represents a single localization event. The format encodes six core observables: the position in the x-dimension, the position in the y-dimension, the acquisition frame index (time of detection), the integrated photon counts of the point spread function (PSF), the channel identifier, and the PSF width estimates in x and y. Additional quality metrics — localization precision, background density, and statistical p-value — are optional but enable the full filtering pipeline when present. The complete column specification is given in Table 2.

All spatial coordinates are expressed in nanometers, and the exact column names listed in Table 2 must be present in the input file; the pipeline validates their presence at load time and skips any file for which a required column is absent. The format does not support 3D (z) coordinates; the pipeline is strictly 2D. In order to process data originating from localization software with differing column conventions, the input should be pre-converted to match the column names specified here before ingestion.

### 4. Pipeline Design

#### 4.1 Modularity

Each stage in the data processing pipeline, such as drift correction, filtering, and clustering, is encapsulated within a module. This ensures that each step can be understood, tested, and modified independently, while the overall workflow remains stable.

#### 4.2 Configuration-driven Behavior

All parameters that can be adjusted within the data processing pipeline have been centralized within a single YAML configuration file. This ensures that there are no thresholds or constants hard-coded within the data pipeline's logic, thus adding reproducibility and batch execution with different configurations.

#### 4.3 Metadata-driven Workflow

Instead of deducing the relevant experimental data from file names and the structure of the folders, this information is kept in a separate Excel file, referred to as the metadata file.

#### 4.4 High-level Structure

The pipeline is orchestrated by a central core that coordinates processing modules. Processing modules transform the data at each step; analysis modules compute quantitative metrics; visualization produces plots and figures; config/utils provide configuration, I/O, and cross-cutting concerns.

##### 4.5 Execution Order and Data Dependencies

For each input file, the pipeline executes the following steps in strict sequence. Later steps depend on the outputs of earlier ones:

*Table 1: Execution Order of Each Modules*

| Step | Module | Description | Data Dependency |
|---|---|---|---|
| 1 | main.py | Load configuration and metadata; discover and iterate over input files | None |
| 2 | pipeline.py | Load raw CSV, validate columns, extract channels, create per-channel files | Raw CSV, metadata |
| 3 | visualization | Initial visualizations and summary metrics | Loaded data |
| 4 | drift_correction | AIM drift correction per channel; align Ch2 to Ch1 drift for dual-channel | Per-channel CSVs |
| 5 | filtering | Sigma, photon count, localization precision, p-value filters | Drift-corrected data |
| 6 | temporal_grouping | Merge repeated localizations into tracks | Filtered data |
| 7 | cell_detection | Optional cell boundary detection | Temporal-grouped data |
| 8 | nearest_neighbor_analysis | KNN distance analysis (optional) | Cell-detected (or grouped) data |
| 9 | clustering | DBSCAN, HDBSCAN, or FINDER clustering per channel | Same as step 9 |
| 11 | clustering | Optional cluster-center KNN | Cluster stats |
| 12 | colocalization | CBC and RE co-localization (dual-channel only) | Same as step 10 |
| 13 | pipeline.py | Export CSVs and images; create stacked histograms; zip outputs | All intermediates |

### 5. Data Flow and Configuration

This section defines the input data format, the metadata-driven workflow for file and channel detection, and the configuration management system. Correct setup of input data and metadata is a prerequisite for the successful execution of a pipeline.

#### 5.1 Input Data Requirements

##### 5.1.1 Format and Column Semantics

The pipeline expects the localization points in the form of CSV files with the following columns:

*Table 2: Description of Columns of Input CSV*

| Column | Semantics | Unit | Mandatory |
|---|---|---|---|
| x (nm) | X-coordinate of localized molecule | nanometers | Yes |
| y (nm) | Y-coordinate of localized molecule | nanometers | Yes |
| z (nm) | Z-coordinate of localized molecule | nanometers | No |
| sigmaX (nm) | Estimated PSF width in x | nanometers | No |
| sigmaY (nm) | Estimated PSF width in y | nanometers | No |
| intensity (photons) | Total photon counts for the localization | photons | No |
| background (photons/nm^2) | Background photon intensity | Photons per $nm^2$ | No |
| p-value | Statistical significance of the localization fit | 0-1 | No |
| localization precision (nm) | Estimated positional uncertainty | nanometers | No |
| channelIndex | Channel identifier (integer) | 0-3 | Yes |
| frameIndex | Frame number during acquisition | integer | Yes |

These columns match the standard output format of dSTORM localization software. The input data must be a 2D localization table with the columns listed in Table 2. The pipeline only works with 2D data and does not support 3D (z) coordinates. The column names must match exactly as specified.

##### 5.1.2 Co-ordinate and Quality Metric Conventions

- Coordinates: All spatial coordinates are in nanometers. The pipeline assumes a flat 2D field of view. The pixel size in the config is used only during drift correction for the transformation between pixel and nm units.

- Localization Precision: Precision is typically computed as $\sigma_{loc} = \sqrt{(\sigma^2 + (a^2/12) + (8\pi\sigma^4 b^2)/(a^2 N))}$, where σ is the PSF width, a is the pixel size, N is the photon count, and b is the background [2]. Here, precision is a measure of error. Higher localization precision means lower localization reliability. Lower localization uncertainty corresponds to higher localization reliability.
- P-value: Lower values indicate stronger evidence that the localization is a real emitter rather than noise. Typical thresholds range from 0.01 to 0.1.

#### 5.2 Metadata-driven Workflow

##### 5.2.1 Purpose of the Metadata File

An Excel metadata file (path: ‘data/metadata/<xlsx_filename>’) drives input file discovery and channel configuration. This design allows:

- Flexible file naming: The pipeline matches input CSVs to metadata rows by base name (without extension). File names need not encode channel information.
- Per-experiment channel mapping: Different experiments may use different channel indices or frame ranges; the metadata specifies these per file.
- Scalability: Adding new experiments or conditions only requires updating the metadata and placing CSVs in the input directory.

##### 5.2.2 Required Metadata Columns

The metadata Excel must have normalized, lowercase, and no-whitespace column names.

- file_name: Base name of the input CSV (e.g., ‘AHA_24014_U2OS-WT_AST0126_AST0063_posXY0_channels_t0_posZ0’)
- first_channel_index: Integer channel index for the first acquired channel
- first_ch_frame_last: Last frame number for channel 1
- second_channel_index: Channel index for the second channel
- second_ch_frame_last: Last frame for channel 2

Optional fields in metadata excel are:

- experiment_nr
- initials
- tag
- 1st channel
- 2nd channel

Channel labels such as AST0063, PST0126 are inferred from the input filename via **extract_channels_from_filename()** and used for filtering and output labeling.

##### 5.2.3 Frame Ranges

The pipeline does not enforce fixed acquisition lengths (e.g., 10,000 or 20,000 frames). Instead, per-file frame ranges are taken from the metadata Excel (last-frame values for channel 1 and channel 2). For each channel, the first skip_frames frames are excluded to account for activation/burn-in at the start of acquisition; this is configured via config/config.yaml under data.input.skip_frames (default: 200). In dual-channel mode, channel 2 is treated as starting immediately after channel 1 ends (i.e., offset by the channel 1 last frame), and the function validates that both channels' frame ranges are positive and ordered correctly.

#### 5.3 Configuration Management

All tunable parameters are available in **config/config.yaml.** The configuration is:

- Loaded once at pipeline startup via **load_config()** in **core/config/loader.py.**
- Validated using Pydantic models in **core/config/models.py** — invalid types or out-of-range values trigger clear errors before any processing begins.
- Passed through to **process_single_file()** and from there to each step function

#### 5.4 Configuration Description

##### 5.4.1 Data Input (data)

| **Key** | **Type** | **Description** | **Example** |
|---|---|---|---|
| data.input.file_format | string | Format of raw localization files | csv (only supported format) |
| data.input.required_columns | list[string] | Column names required in input CSV | "x (nm)", "y (nm)", "channelIndex", "frameIndex" |
| data.input.xlsx_filename | string | Metadata Excel filename (placed in data/metadata/) | "dSTORM Data_Input.xlsx" |
| data.input.skip_frames | number | Number of frames to skip at the start of each channel's frame range | e.g. 200 |
| data.input.required_metadata_columns | list[string] | Column names required in metadata xlsx | file_name, first_ch_frame_last, second_channel_index, second_ch_frame_last |

##### 5.4.2 Channel Display (channels)

| Key | Type | Description | Example |
|---|---|---|---|
| channels.Ch1.color | string | Hex color for Channel 1 | "#FF00FF" |
| channels.Ch2.color | string | Hex color for Channel 2 | "#00FFFF" |

##### 5.4.3 Visualization

| Key | Type | Description | Example |
|---|---|---|---|
| visualization.use_napari | boolean | Whether to save napari plots or not. | e.g. true or false |

##### 5.4.4 Filtering

| Key | Type | Description | Example |
|---|---|---|---|
| filtering.intensity.min_value | number | Intensity threshold value | e.g. 200 |
| filtering.localization_precision.threshold_value | float | Max allowed localization precision (nm) | e.g. 20.0 |
| filtering.p_value.threshold_value | float | Max allowed p-value | 0–1 |
| filtering.sigma.min_value | float | Min allowed σ (nm) | e.g. 50.0 |
| filtering.sigma.max_value | float | Max allowed σ (nm) | >min_value |

##### 5.4.5 Drift Correction (drift_correction)

| Key | Type | Description | Example |
|---|---|---|---|
| drift_correction.pixel_size | number | Pixel size (nm) | e.g. 100 |
| drift_correction.segmentation | integer | Temporal segments | e.g. 20 |
| drift_correction.intersect_d_nm | float | Intersection distance (nm) | e.g. 60 |
| drift_correction.roi_r_nm | float | ROI radius (nm) | true or false |
| drift_correction.drift_validation.enable | boolean | Enable drift validation | e.g. 15.0 |
| drift_correction.drift_validation.max_segment_drift_nm | float | Max allowed segment drift (nm) | e.g. 10 |
| drift_correction.drift_validation.n_previous_segments | integer | Previous segments used for replacement | true or false |

| | | | |
|---|---|---|---|
| drift_correction.sanity_checks.filter_boundary | boolean | Remove x ≤ 0 or y ≤ 0 | true or false |
| drift_correction.sanity_checks.filter_parameters | boolean | Remove sx ≤ 0, sy ≤ 0, lp ≤ 0 | true or false |

##### 5.4.6 Temporal Grouping (temporal_grouping)

| Key | Type | Description | Options |
|---|---|---|---|
| temporal_grouping.use | boolean | Enable temporal grouping | true/false |
| temporal_grouping.max_frame_gap | integer | Max frame gap | e.g. 2 |
| temporal_grouping.max_distance_nm | float | Max linking distance (nm) | e.g. 50 |
| temporal_grouping.min_duration | integer | Min track duration | e.g. 1 |
| temporal_grouping.max_duration | integer | Max track duration | ≥ min_duration |

##### 5.4.7 Cell Detection (cell_detection)

| Key | Type | Description | Options |
|---|---|---|---|
| cell_detection.use | boolean | Enable cell detection | true/false |
| cell_detection.approach | string | Algorithm | "grid" or "vornoi" |
| cell_detection.grid.grid_size | integer | Grid bins per axis | e.g. 100 |
| cell_detection.grid.high_density_threshold | integer | Min localizations per bin | e.g. 3 |
| cell_detection.vornoi.percentile | float | Percentile before Otsu threshold | 1–100 (e.g. 99.0) |

##### 5.4.8 Nearest Neighbor Analysis (nearest_neighbor_analysis)

| Key | Type | Description | Options |
|---|---|---|---|
| nearest_neighbor_analysis.use | boolean | Enable NN analysis | true/false |
| nearest_neighbor_analysis.radius | float | Radius (nm) | e.g. 100 |
| nearest_neighbor_analysis.k | integer | Number of neighbors | e.g. 1 |
| nearest_neighbor_analysis.algorithm | string | sklearn algorithm | "auto", "ball_tree", "kd_tree", "brute" |
| nearest_neighbor_analysis.metric | string | Distance metric | "euclidean" |

5.4.9 Clustering

| Key | Type | Description | Options |
|---|---|---|---|
| clustering.use | boolean | Enable clustering | true/false |
| clustering.use_cluster_knn | boolean | Enable cluster centroid KNN | true/false |
| clustering.ch1.use | boolean | Whether to run clustering on ch1 or not | true/false |
| clustering.ch1.method | string | Clustering method (Ch1) | "dbscan", "hdbscan", "finder" |
| clustering.ch1.dbscan.eps | float | DBSCAN radius (Ch1) | e.g. 100 |
| clustering.ch1.dbscan.min_samples | integer | DBSCAN min samples (Ch1) | e.g. 10 |
| clustering.ch1.hdbscan.min_cluster_size | float | Minimum size of cluster | e.g. 10 |
| clustering.ch1.hdbscan.min_samples | integer | HDBSCAN min samples (Ch1) | e.g. 10 |
| clustering.ch1.finder.threshold | integer | Minimum points for cluster | e.g. 10 |
| clustering.ch1.finder.points_per_dimension | integer | Grid resolution for parameter search | e.g. 15 |
| clustering.ch1.finder.algorithm | list[string] | Inner algorithm | e.g. "dbscan", "DbscanLoop" |
| clustering.ch1.finder.min_threshold | integer | Minimum threshold to search | e.g. 5 |
| clustering.ch1.finder.max_threshold | integer | Maximum threshold to search | e.g. 21 |
| clustering.ch1.finder.decay | float | Decay for parameter search | e.g 0.5 |
| clustering.ch2.use | boolean | Whether to run clustering on ch2 or not | true/false |
| clustering.ch2.method | string | Clustering method (Ch2) | Same as ch1 |
| clustering.ch2.dbscan.eps | float | DBSCAN radius (Ch2) | e.g. 100 |
| clustering.ch2.dbscan.min_samples | integer | DBSCAN min samples (Ch2) | e.g. 10 |
| clustering.ch2.hdbscan.min_cluster_size | float | Minimum size of cluster | e.g. 10 |
| clustering.ch2.hdbscan.min_samples | integer | HDBSCAN min samples (Ch1) | e.g. 10 |

| | | | |
|---|---|---|---|
| clustering.ch2.finder.threshold | integer | Minimum points for cluster | e.g. 10 |
| clustering.ch2.finder.points_per_dimension | integer | Grid resolution for parameter search | e.g. 15 |
| clustering.ch2.finder.algorithm | list[string] | Inner algorithm | e.g. "dbscan", "DbscanLoop" |
| clustering.ch2.finder.min_threshold | integer | Minimum threshold to search | e.g. 5 |
| clustering.ch2.finder.max_threshold | integer | Maximum threshold to search | e.g. 21 |
| clustering.ch2.finder.decay | float | Decay for parameter search | e.g 0.5 |

5.4.10 Co-localization

| Key | Type | Description | Options |
|---|---|---|---|
| colocalization.cbc.radius | float | CBC radius (nm) | e.g. 100 |
| colocalization.cbc.n_steps | integer | Radius steps | e.g. 10 |

### 6. Implementation

This section provides a detailed description of each processing step: the algorithms used, the reason for design choices, key parameters, and expected outputs.

#### 6.1 Data Loading and Channel Extraction

6.1.1 What is done?

- Load full CSV via load_data() from utils/data_handling.py. The raw CSV contains localizations co-ordinates from all channels and frames.
- Validate required columns: If any column from config.data.input.required_columns is missing, the pipeline logs an error and skips the file.
- Look up metadata: The metadata row for this file is retrieved by matching the base name to the file name column.
- The code filters localisations for each channel by selecting rows where channelIndex matches and frameIndex is within a specific range, from the channel's start frame plus skip_frames (default 200, set in config.yaml) up to the last frame for that channel. This produces separate DataFrames for each channel.

- Save per-channel CSVs: Each channel subset is written to *_ch1.csv and *_ch2.csv in the input directory. These temporary files are used by the AIM drift correction module, which expects a single-channel CSV per run.
- Combine channels: For downstream steps, channels are concatenated into a single Data Frame. The channelIndex column is preserved so that each step can operate per channel when needed.

###### 6.1.2 Why this designed?

Drift correction must be applied independently to each channel because the two channels are acquired sequentially. Each channel experiences different drift during its acquisition window. After correcting each channel's drift, the pipeline aligns channel 2 to channel 1 by subtracting the cumulative drift of channel 1 from all channel 2 coordinates. This ensures that both channels are registered to a common coordinate system for co-localization and inter-channel analyses.

#### 6.2 Drift Correction: Adaptive Intersection Maximization (AIM)

Due to mechanical drift of the stage or sample during data acquisition, localisations shift over time. If uncorrected, this drift can blur the reconstructed image and negatively affect spatial analysis. AIM corrects this drift by identifying stable reference points—localisations that appear in multiple temporal segments—and estimating the displacement between these segments. The method does not require fiducial markers or separate reference channels. The implementation used here is adapted from the AIM method implemented in Picasso [3] and modified for pipeline integration, including CSV input/output handling, sanity checks, drift validation, and HDF5 conversion for internal processing.

###### 6.2.1 Algorithm

1. **Temporal Segmentation**: The acquisition is divided into N equal segments (default: N = 100). Each segment contains a subset of frames.
2. **Reference Selection**: In the first round, the first segment is used as the reference. In the second round, the entire (partially corrected) dataset is used as a reference to refine the first segment as well.
3. **Per-segment Drift Estimation:**
    - Convert reference and target localizations to a discretized grid in units of "intersect_d" (intersection distance).
    - For each shift within a local search region (ROI) of size "roi_r," calculate the number of overlapping reference and target localizations (occupying the same grid cell).
    - The shift that maximizes this overlap is defined as the coarse drift estimate.
    - Apply FFT-based sub-pixel refinement to obtain a precise drift estimate.

4. **Interpolation:** Cubic spline interpolation (with boundary extension) is used to obtain per-frame drift from the per-segment drift estimates.
5. **Coordinate correction:** For each localization, subtract the drift corresponding to its frame index.
6. **Two-round refinement:** The first round corrects drift relative to the initial segment; the second round uses the full dataset as a reference, improving accuracy for the early frames and reducing bias.

###### 6.2.2 Key Parameters

- pixel_size: Camera pixel size in nm. Used to convert between pixel and nm units internally. Must match the acquisition setup.
- segmentation: Number of temporal segments. More segments yield finer temporal resolution but increase noise; fewer segments are smoother but may miss rapid drift.
- intersect_d_nm: Distance (in nm) used for discretising coordinates when counting overlaps. Should be on the order of localisation precision (e.g., 20 nm).
- roi_r_nm: Radius of the local search region in nm. Must be larger than the expected drift per segment. Too small underestimates drift; too large increases computation and can introduce noise.

###### 6.2.3 Drift Validation

It is a newly added component to the existing implementation. When enabled, segments with drift magnitude exceeding max_segment_drift_nm are flagged as outliers. These are replaced by the mean of the previous N valid segments. This mitigates artefacts from sparse segments, blinking, or acquisition glitches that can produce unrealistic drift spikes.

###### 6.2.4 Sanity Checks

After drift correction, localizations that would fall outside valid bounds or have invalid parameters are removed:

- filter_boundary: Drop localizations with x≤0 or y≤0 (outside image)
- filter_parameters: Drop localizations with sx≤0, sy≤0, or lp≤0

Removed localizations are logged; the pipeline continues with the remaining data.

#### 6.3 Filtering

Filters are applied step-by-step in a chosen order to remove the most problematic localisations first. Each filter step reduces the dataset; downstream filters operate on the already-filtered result. The applied filters are described below.

##### 6.3.1 Sigma Filter

PSF width (σ) represents the quality of the Gaussian fit. Values far from the expected PSF size indicate misfits, de-focus, or non-Gaussian artefacts. The Sigma filter removes localisations with poor PSF fits.
**Criterion**: sigma_min ≤ σx, σy ≤ sigma_max

##### 6.3.2 Photon Count Filter

Removes noisy localisations by keeping only those with photon counts above a fixed minimum value, regardless of channel type.

##### 6.3.3 Localization Precision Filter

The localisation precision refers to the certainty level that we have about the location of a single emitter. This filter will eliminate localisation whose precision is greater than the defined threshold.

##### 6.3.4 P-value Filter

A low p-value indicates strong evidence that the detection is a real emitter. This filter removes statistically unreliable localisations. It keeps localisations with p-value below the threshold.

#### 6.4 Temporal Grouping

In direct Stochastic Optical Reconstruction Microscopy (dSTORM), temporal grouping refers to the post-processing step of merging multiple, sequential localised detections (blinks) of the same fluorophore into a single, accurate coordinate [4]. Without temporal grouping, the same molecule contributes multiple localisations, artificially inflating density and cluster sizes. The temporal grouping method performs spatial-temporal clustering using fixed thresholds for both spatial distance and frame separation [5]. This approach does not adapt to localisation precision and instead applies globally defined constraints.

##### 6.4.1 Methodology

1. **Spatial Indexing:** A cKDTree is constructed over all localization coordinates to enable efficient neighbor queries.
2. **Spatiotemporal Clustering:** For each unvisited localization, a breadth-first search is performed to identify neighboring detections that satisfy both:
    - Spatial proximity: within a fixed distance threshold
    - Temporal proximity: within a predefined maximum frame gap

   All detections connected under these constraints form a single group.
3. **Group Merging:** Each group is merged into a single representative localization using the pooling strategy:

    - Mean spatial co-ordinates
    - Summed photon counts
    - Aggregated precision and auxiliary parameters
4. **Duration-based filtering:** Groups can be filtered based on their temporal duration:
    - Exclude short-lived tracks (e.g., noise or transient artifacts)
    - Remove anomalously long tracks

###### 6.4.2 Parameters

- max_frame_gap: This is the largest allowed frame difference between two candidate localizations to still consider them part of the same track.
- max_distance_nm: This is the fixed spatial radius used to decide whether two localizations are close enough to be linked.
- max_duration: Tracks longer than this are removed. This helps reject suspiciously long tracks that might be coming from over-linking or artefacts.
- min_duration: After linking, each track gets a duration measured in frames. Tracks shorter than this are removed.

#### 6.5 Cell Detection (Optional)

Cell detection limits analysis to localisations that lie within cellular regions, reducing the influence of extracellular or non-cellular background. This step is optional and can be skipped so that the entire field of view is used for clustering and co-localization.

###### 6.5.1 Grid-based Cell Detection

**Methodology**

1. **Binning:** The field of view is divided into a 2D grid. The number of bins along each axis is set by grid_size, so the grid has grid_size × grid_size cells. Bin edges are defined by linearly spacing between the minimum and maximum x and y in the dataset.
2. **Assignment**: Each localization is assigned to a bin via digitization; columns x_bin and y_bin is added. Only rows with valid bin indices (0 to grid_size - 1) are kept.
3. **Density:** For each bin, the number of localizations is counted. Bins with count ≥ high_density_threshold are classified as high-density (cell region).
4. **Filtering:** A boolean column is_cell is set to True for localizations in high-density bins. The output Data-frame contains only rows with is_cell == True.

**Parameters**

- grid_size: Number of bins per axis. Larger values give finer spatial resolution but smaller counts per bin; smaller values give coarser regions.
- high_density_threshold: Minimum localizations per bin to consider the bin as part of the cell. Bins below this are treated as background.

##### 6.5.2 Voronoi-based Cell Detection

**Methodology**

1. **Voronoi tessellation:** A Voronoi diagram is computed from all localization coordinates (x, y). Each point has an associated Voronoi cell (region of space closer to that point than to any other).
2. **Cell area:** For each finite Voronoi region, the polygon area is computed using the shoelace formula on the region vertices. Infinite or empty regions are excluded. The area is stored in a new column voronoi_area.
3. **Clipping:** To reduce the influence of extreme, often boundary, regions, Voronoi areas are clipped at the clip_percentile (e.g. 99th percentile). Only finite areas ≤ this clip value is used for thresholding.
4. **Thresholding:** Otsu's method is applied to the clipped Voronoi areas to automatically choose a threshold that separates two classes. Points whose Voronoi area is below (or equal to) this threshold are classified as "cell region"; the rest as background.
5. **Filtering:** A column is_cell is set accordingly. The returned Data-frame contains only localizations with is_cell == True. The Voronoi object is also returned for optional visualization.

**Parameters**

- clip_percentile: Percentile used to clip the largest Voronoi areas before Otsu thresholding. Reduces the impact of edge and outlier regions.

#### 6.6 Nearest Neighbor Analysis (Optional)

Nearest neighbor analysis quantifies spatial relationships between localizations, either within a single channel (intra-channel) or between two channels (inter-channel). It supports localization-level characterization (e.g. distance distributions) and is used again at the cluster level when cluster-center KNN is enabled.

##### 6.6.1 Methodology

- **Configuration:** The approach is read from config. Parameters k, algorithm, metric, and radius are used for nearest neighbor and mean-distance-within-radius computations.

- **Intra-channel Analysis:** For each channel (Channel1, Channel2), localizations are subset by channelIndex. For each point, the k nearest neighbors within the same channel are found. Distances are stored; the first nearest neighbor distance is used for histograms and a new column.
- **Inter-channel Analysis:** For each ordered pair of distinct channels (Ch1→Ch2, Ch2→Ch1), distances from each localization in the "primary" channel to the nearest localization in the "reference" channel are computed. New columns are added; histograms and CSVs are saved.
- **Mean distance within Radius:** For a fixed radius r, the mean distance from each point to all neighbors within radius r is computed (intra and, for two channels, inter). Results are stored in columns, with corresponding histograms and CSVs.

###### 6.6.2 Parameters

- use: whether to run this step or not in the pipeline. e.g. true/false
- k: Number of nearest neighbors to compute (e.g., $k = 1$ for the first nearest neighbor).
- algorithm: Search algorithm used by NearestNeighbors. Possible options: auto, ball_tree, kd_tree, or brute.
- metric — Distance metric used for neighbor computation (e.g., Euclidean).
- radius: Radius (in nanometers) used for computing the mean distance of neighbors within a specified spatial radius.

#### 6.7 Clustering

Spatial clustering groups localizations into molecular groups (e.g. protein clusters). The pipeline supports three methods per channel: DBSCAN, HDBSCAN, and FINDER. Each channel can use a different method. Clustering is run on the same data used for optional cell detection and optional background filter.

###### 6.7.1 Methodology

- Per-channel Processing: For each channel (Ch1, Ch2), localizations are filtered by channelIndex. Coordinates are taken from the x and y columns.
- Method Call:
    - DBSCAN: sklearn.cluster.DBSCAN is fitted with eps (nm) and min_samples. Points not in any cluster receive label -1.
    - HDBSCAN: sklearn.cluster.HDBSCAN is fitted with min_cluster_size, min_samples, and cluster_selection_epsilon. Again, noise is labelled -1.
    - FINDER: The C++ extension finder_cpp.run_finder_2d is called with the channel's finder parameters. FINDER explores a range of (ε, minPts)-like parameters and returns optimal labels plus chosen parameters (e.g. sigma, threshold). These are saved to a JSON file per channel.
- Label Offset: For dual-channel data, cluster IDs of Ch2 are offset so they do not collide with Ch1. Noise remains -1.

- Assignment: The combined DataFrame receives a cluster column. Each row is assigned the cluster label of its channel.
- Cluster Statistics: For each channel, save_cluster_stats() builds a per-cluster table: cluster ID, channel index, channel label, n_points, centroid (x_center, y_center), standard deviations, convex-hull area, and density. Clusters with fewer than three points get an area of 0. A row for "noise" (cluster -1) is appended. The table is saved.
- Outputs: The clustered DataFrame is saved. Cluster stat CSVs and JSON (for FINDER) are written to the clustering output directory. Downstream, histograms of cluster size and area are generated, and summary statistics are appended for reporting.

###### 6.7.2 DBSCAN

DBSCAN (Density-Based Spatial Clustering of Applications with Noise) is a density-based clustering algorithm designed to discover clusters of arbitrary shape in spatial databases [6].

**Parameters**

- **eps**: Maximum distance (nm) between two points for them to be in the same neighborhood. Critical for cluster extent.
- **min_samples**: Minimum points in a neighborhood for a core point. Smaller values yield more, smaller clusters; larger values favor fewer, denser clusters.

###### 6.7.3 HDBSCAN

HDBSCAN (Hierarchical Density-Based Spatial Clustering of Applications with Noise) is a density-based clustering algorithm that builds a hierarchy of clusters based on varying data density and extracts the most stable clusters while identifying outliers [7].

**Parameters**

- **min_cluster_size**: Minimum number of points in a cluster.
- **min_samples** : Used for core distance / stability.

###### 6.7.4 FINDER

FINDER is an unbiased clustering algorithm for single-molecule localization microscopy that automatically selects global parameters to robustly identify molecular clusters while minimizing false positives and false negatives in noisy and densely populated data [8].

**Algorithm**

1. **Parameter Search:** A grid of (threshold, sigma)-like parameters are defined (e.g. min_threshold to max_threshold, with points_per_dimension controlling resolution).
2. **Clustering:** For each parameter combination, DBSCAN is run. Cluster labels are obtained.
3. **Similarity score:** A similarity score (e.g. based on silhouette or parameter sensitivity) is computed for each combination.
4. **Selection:** The combination that maximizes the score (with optional decay for robustness) is chosen. The corresponding labels and chosen parameters are returned.

**Parameters**

- threshold: This is the baseline DBSCAN minPts-like value (minimum neighbors to treat a point as dense/core).
- points_per_dimension: This controls how finely FINDER samples the parameter space.
- algorithm: This picks the clustering backend used for each tested parameter pair, e.g. "dbscan" or "DbscanLoop".
- min_threshold, max_threshold: Range of the threshold parameter in the search.
- decay: This is the selection sensitivity for choosing the final point on the score-optima curve.

#### 6.8 KNN Analysis of Cluster Centroids (Optional)

When clustering.use_cluster_knn is true, a second pass of nearest-neighbor analysis is run on cluster statistics (centroids and areas), not on raw localizations. This characterizes relationships between clusters. For example, the distance from a first channel cluster to the nearest second channel cluster.

##### 6.8.1 Methodology

- Input: Cluster statistics Data-frames (one per channel), with noise (cluster -1) removed and cluster IDs prefixed by channel (e.g. ch0_0, ch1_0) to avoid collisions. These are concatenated into a single table with columns including x_center, y_center, area_nm2, and cluster.
- Cluster KNN: analyze_nearest_neighbors() is called with this combined stats table, using x_col='x_center', y_col='y_center', and is_clustered=True. Intra- and inter-channel distances between cluster centroids are computed (and optionally mean within

radius). Histograms and CSVs are saved with a cluster_ prefix; results are merged into the nearest-cluster DataFrame (e.g. columns nn_distance_to_<other_channel_label>_nm).

- All plots are saved in the clustering output directory. Summary statistics from cluster KNN are appended to the pipeline's summary for reporting.

#### 6.9 Co-localization Analysis

Co-localization quantifies the spatial association between two molecular species (two channels). The pipeline implements two complementary methods: Coordinate-Based Co-localization (CBC) and Relative Enrichment (RE). This step runs only when two channels are present, and colocalization use is true.

##### 6.9.1 Co-ordinate-Based Co-localization (CBC)

Purpose: For each localization in one channel, measure the distance to the nearest localization in the other channel. The fraction of points within a given radius summarizes spatial overlap [10].

**Methodology**

1. **Data preparation:** The combined localization DataFrame is split by channelIndex into two channel Data-frames. Columns are renamed to Locan conventions (position_x, position_y). Each subset is converted to a locan.LocData object.
2. **Bidirectional CBC:** CBC is computed in both directions: Ch1→Ch2 (primary Ch1, reference Ch2) and Ch2→Ch1 (primary Ch2, reference Ch1). For each direction, Locan's CoordinateBasedColocalization is called with the chosen radius and n_steps (number of radius steps for a sweep).
3. **Output:** For each direction, the primary channel's coordinates and the resulting co-localization values are stored. These are used for plotting (e.g. scatter or radius curves) and summary statistics.

**Parameters**

- **Radius**: This is the maximum spatial scale considered for CBC. Think of it as: "Up to what distance should I consider two channels potentially co-located?"
- **n_steps**: This sets how many intermediate radius values are evaluated between 0 and radius.

##### 6.9.2 Relative Enrichment (RE)

Purpose: Quantify whether the primary channel is enriched or depleted relative to the reference channel in local neighbourhoods defined by the reference channel's Voronoi tessellation, compared to a random expectation [9].

**Methodology**

1. **Voronoi tessellation:** In relative_enrichment(), the **reference** channel's localizations are used to build a 2D Voronoi diagram. Each reference point has a Voronoi cell (region). Only finite regions are kept; a boolean index marks valid cells.

2. **Primary points per region:** For each valid reference cell, the number of **primary** channel localizations falling inside that cell is counted (using polygon containment and a cKDTree to limit the primary points queried). This gives, per reference localization, a count n_points of primary localizations in its neighborhood, the cell **area**, and the **first-order mean distance** to neighboring reference points.

3. **Relative enrichment per localization:** In compute_relative_enrichment(), an expected count is derived from the total number of primary points and the ratio of (region area) to (total area). The ratio of observed to expected (or a similar normalization) gives a per-reference-localization RE value. Values > 1 indicate enrichment of the primary channel near that reference point; < 1 indicate depletion.

#### 6.10 Summary Statistics and Reporting

Throughout the pipeline, summary statistics are collected and appended to shared output structures, allowing a single experiment run to be summarized in tables or downstream scripts.

- Processing summary: After each major step (initial, after_drift_correction, after_filtering, after_temporal_grouping, after_background_filter, after_cell_detection), metrics (e.g. counts, basic stats) are gathered by plot_metrics() and passed to append_processing_summary_stats(). These are written under the output directory so that retention rates and step-wise metrics are available.
- Clustering summary: Histogram and cluster-stat plot results are passed to append_clustering_summary_stats() and saved in the same output area.
- Nearest neighbour summary: Results from localisation-level nearest neighbour and (when enabled) cluster-level KNN are passed to append_nn_summary_stats().
- Co-localisation summary: CBC and RE summary data frames or statistics are passed to append_coloc_summary_stats().
- Exact file names and formats are defined in utils/utils.py and the pipeline; they aggregate per-step and per-channel information for easy comparison across experiments or conditions.

#### 6.11 Export of Results

The final step consolidates all intermediate data and figures into the experiment's output folder and creates a single archive for sharing or archiving.

##### 6.11.1 What is done?

- Stacked histograms: For each of the metrics lp, bg, sx, sy, and pvalue, create_stacked_histogram() is called. It looks for the step-specific histogram images (initial, after_drift_correction, after_sigma_filter, after_photons_count_filter, after_localization_precision_filter, after_pvalue_filter, after_temporal_grouping, after_cell_detection) in the temporary directory structure. For each metric, it stacks these images vertically into a single image and saves it as e.g. {metric}_hist_all_steps.png in the temp tree. Only the steps that produced a file are included.
- Consolidation of Files
    - All CSV files from the temp tree are moved into temp/test_data/ via move_all_files().
    - CSVs from the sub-folders drift_corrected, filtered, temporal_grouped, cell_detected, and from quantification_results/clustering and quantification_results/distance_based are moved into temp/test_data/.
    - All images (e.g. .png) under the temp tree are moved into temp/test_images/ via move_all_images(), including images from quantification_results/clustering, quantification_results/distance_based, and quantification_results/colocalization.

- ZIP archive: The entire temp directory is zipped using zip_and_rename_folder(). The archive name is the experiment base name (e.g. derived from the input file name), and the archive is written to data/output/. If a file with that name already exists, a unique name is generated (suffix with a letter or number) to avoid overwriting.
- After export, the pipeline considers the run complete for that file. The temp directory is later cleared by the main entry point when all files have been processed.

#### 6.12 Output Layout (per experiment)

- data/output/*.zip
- .zip contains:
    - test_data/ — All CSV outputs (initial, drift-corrected, filtered, temporal-grouped, cell-detected, clustering, distance-based, co-localization, and any summary CSVs moved there).
    - test_images/ — All images: step-wise histograms, stacked histograms, cluster and NN plots, CBC and RE figures, cell detection and Voronoi images, etc.

- Unzipping the archive reproduces the full set of data and figures for that experiment in a portable structure.

#### 7. Post-processing Scripts

##### 7.1 merge-csv

It is the batch CSV merger. It scans the output folders, groups files by cell line and channel, normalizes channel-name variants, and writes merged summary CSVs. It has separate paths for cluster stats, intra-channel results, inter-channel results, and mean-distance outputs, including the inter-channel mean-distance case. The main job is to consolidate many per-run CSVs into cleaner combined files.

##### 7.2 save-napari-visualization

It automates figure generation. It loads the project's config and metadata, finds which channels belong to each dataset, opens the stepwise CSV files in each output folder, saves Napari screenshots for each processing stage, stacks them into a composite image, and also creates cluster visualizations when clustering data exists. In short, it is the "export all visual outputs" script.

##### 7.3 visualization-steps

It is the step-by-step viewer. It looks in the configured output folder, lists the available pipeline stages for the current experiment, lets you pick one from a terminal menu, and then opens that stage in Napari. It is a lightweight way to inspect Initial, Drift Correction, Filtering, Temporal Grouping, and Cell Detection outputs one at a time.
